# Length and time scales of cell-cell signaling circuits in agar

**DOI:** 10.1101/220244

**Authors:** Joy Doong, James Parkin, Richard M. Murray

## Abstract

A community of genetically heterogeneous cells embedded in an unmixed medium allows for sophisticated operations by retaining spatial differentiation and coordinating division-of-labor. To establish the principles of engineering reliable cell-cell communication in a heterogeneous environment, we examined how circuit parameters and spatial placement affect the range of length and time scales over which simple communication circuits interact. We constructed several “sender” and “receiver” strains with quorum-sensing signaling circuits. The sender cell colony produces acyl homoserine lactones (AHL), which diffuse across the semisolid medium. The receiver cell colony detects these signal molecules and reports by fluorescence. We have found that a single colony of one sender variant is sufficient to induce receiver response at more than 1.5cm separation. Furthermore, AHL degradase expression in receiver colonies produces a signal threshold effect and reduces the response level in subsequent receiver colonies. Finally, our investigation on the spatial placement of colonies gave rise to the design of a multicellular long-range communication array consisting of two alternating colony types. Its signal response successfully propagated colony-by-colony along a six-colony array spanning 4.8cm at a transmission velocity of 12.8 hours per colony or 0.075cm per hour. In addition, we have developed a reaction-diffusion model that recreates the observed behaviors of the many performed experiments using data-informed parameter estimates of signal diffusion, gene expression, and nutrient consumption. These results demonstrate that a mixed community of colonies can enable new patterning programs, and the corresponding model will facilitate the rational design of complex communication networks.

## Introduction

Research in synthetic biology involves identifying effective algorithms and strategies in living systems and applying this knowledge to artificial biological design. This approach has proven successful for small genetic circuits in uniform cultures, such as Boolean logic networks and oscillators in bacteria [9, 17], but to perform more complicated tasks will require advanced strategies.

One emerging strategy is microbial consortia, the design of multiple, genetically distinct strains of cooperating microbes to achieve a desired function [5, 15]. A mixed community of engineered cells can perform more complex functions than a homogeneous population because dividing a complex pathway between genetically distinct,cooperativepopulationsalleviatestherobustness-complexitytradeoff associated with synthetic gene circuits [3, 16]. The importance of coordination to microbial communities cannot be understated. From division-of-labor metabolic relationships to opportunistic human pathogens that simultaneously attack compromised tissues [12, 13], nearly all microbes coordinate within the species and between the species boundaries. Establishing the principles to engineer reliable coordinated behaviors is therefore an important step towards more advanced synthetic biology applications.

Recent studies have described engineered synthetic consortia capable of a range of coordinated behaviors, such as ecological dynamics [18], population dynamics [16] and pattern formation [3, 4, 7, 11]. Some of these works have applied multiple microbial strains to semisolid media preparations [3]. The spatial heterogeneity of these media has enabled the band-detect behavior [3] and could permit other new patterning interactions and networks. However, the range of possible communication distance and time scale in semisolid media has not been explicitly explored. This information will help outline the range of patterning behaviors that are experimentally achievable from simple interactions and coordination, providing the foundation for future research into more complex patterning programs.

Here, we sought to investigate the engineering principles for reliable cell-cell communication in centimeter-scale heterogeneous environments. We examined how circuit parameters and spatial placement affect the range of length and time scales over which simple communication circuits interact in a semisolid medium. These signaling circuits utilize quorum-sensing (QS) components to program “sender” and “receiver” strains (Figure 1). The sender cell colony produces acyl homoserine lactone (AHL) signal molecules, which diffuse across semisolid media. The receiver cell colony detects the AHL signals and reports by fluorescence. We tuned the signaling radius, the maximum distance separating a sender colony from a receiver colony it can communicate with, by varying the ribosome binding site (RBS) strength of either the sender or receiver circuit. We have found that a single colony of one sender strain variant is sufficient to induce receiver response at more than 1.5cm separation. Furthermore, we demonstrated that AHL degradase expression in receiver colonies produces a signal threshold effect and reduces the response level in subsequent receiver colonies.

**Figure 1:**
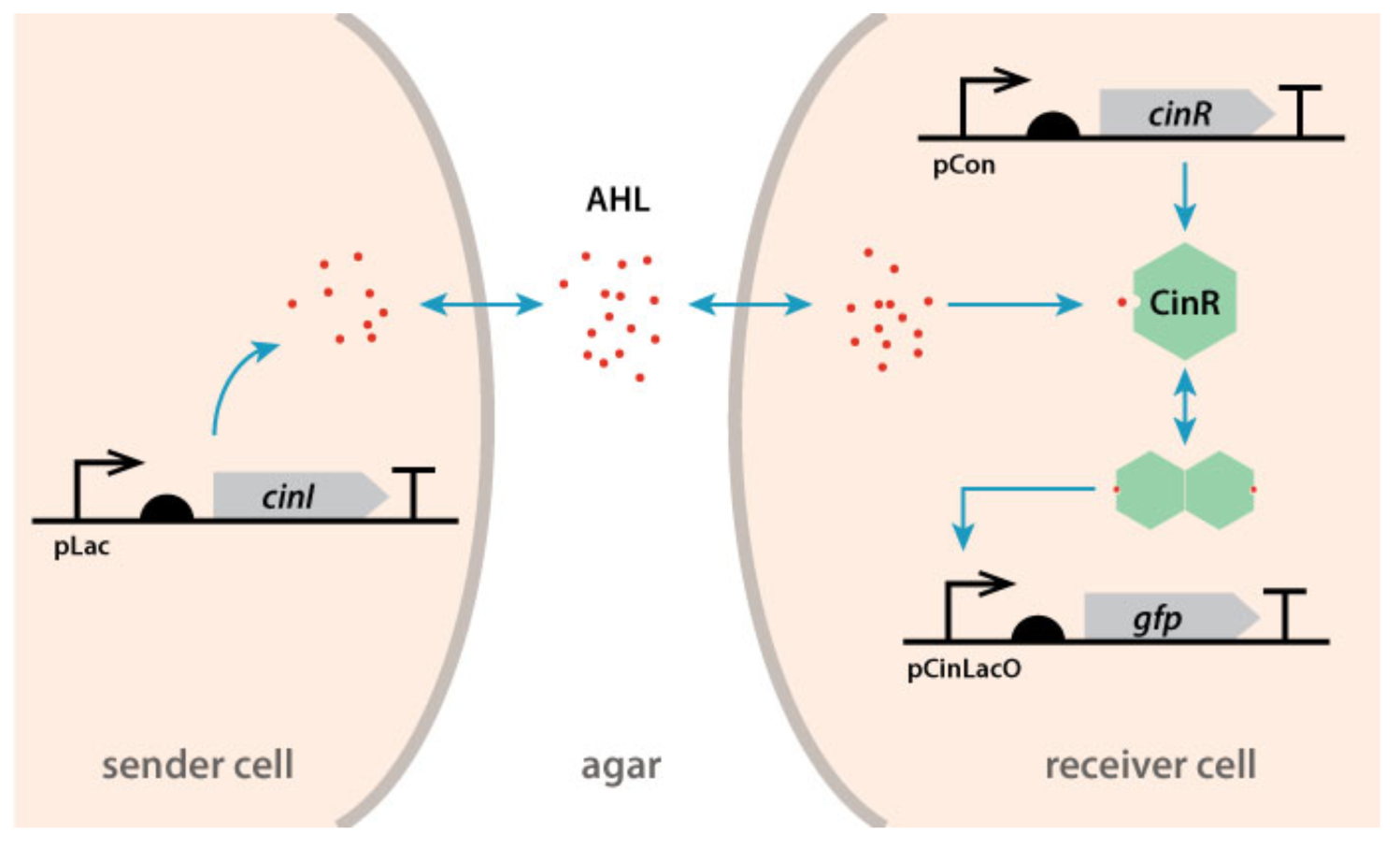
Example of a pair of sender-receiver signaling circuits assembled. In the sender cell, the synthase CinI produces one type of signal AHL, which diffuses across agar. In the receiver cell, AHL binds CinR to form a complex, which activates the expression of GFP.

Building on the sender-receiver paradigm, we sought to achieve longer signaling distances by establishing a communication cascade in which the AHL emitted from sender colonies initiates signaling from neighboring sender colonies. We envisioned connecting distinct colonies in a communication network consisting of two orthogonal chemical channels based on spatial proximity and separation. Tamsir et al. have formerly demonstrated a similar colony-based system that was ‘wired’ with QS-based cell-cell communication [19]. However, the relative spatial orientation of their colonies is not relevant to the function of the device because the communication channels are differentiated solely by orthogonal chemical signals and not by spatial separation. In our work, we demonstrate a system where an intentional spatial pattern is critical for the function of the system. This system consists of an array of two alternating colony types for the purpose long-range communication (Figure 2). Each colony in this system communicates with immediately adjacent colony to propagate the signal colony by colony. Hence, only two orthogonal chemical channels are required for this six-colony propagation array. This resultant ‘wiring’ structure can be expanded to larger signaling networks and patterns in future work.

**Figure 2:**
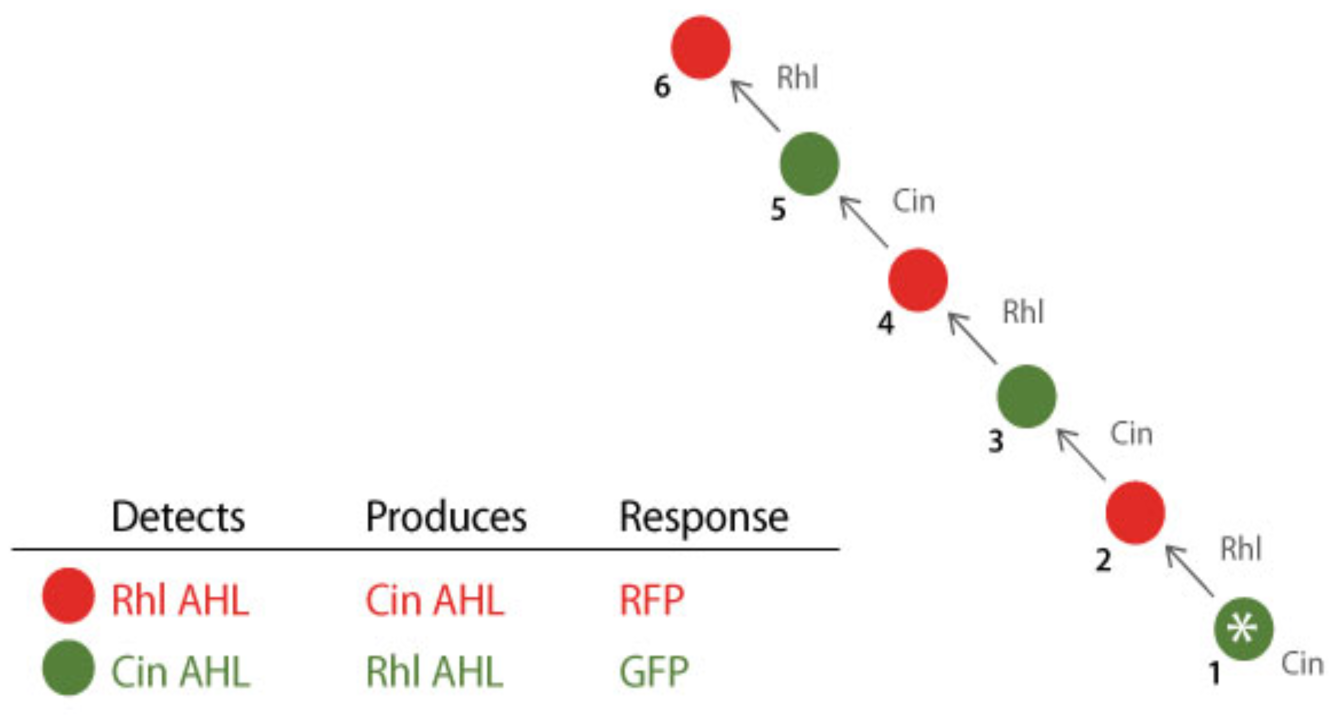
The signal-conducting array consists of two alternating strains, such that the signal detected and sent are orthogonal in each colony. Rhl AHL: N-butyryl-L-homoserine lactone (C_4_-HSL) Cin AHL: N-(3-Oxotetradecanoyl)-L-homoserine lactone (3-Oxo-C14-HSL)

## Results

### Quorum sensing communication circuit produces a range of useful response distance in semisolid media

We first characterized the range of response distance and time that our QS communication circuits could achieve by testing different arrangements of two sender strain variants and four receiver strain variants in Luria Bertani (LB) agar plates. Three of the receiver strains were transformed with circuits that differed only in the strength of the RBS (Figure 3) upstream of the green fluorescent protein (GFP) reporter and of the QS transcription factor. The comparison of these receiver strains demonstrated that exchanging RBS significantly altered the signaling radius of the response, as seen in Figure 3. In this experiment, a row of strong RBS variant sender colonies was seeded above lines of receiver colonies. Using a weak RBS upstream of the transcription factor remarkably increased the basal GFP level and reduced the response by the circuit. The effect was sufficiently remarkable to render this variant useless. On the other hand, alternate RBSs upstream of the GFP marker can dramatically impact the signaling radius.

In order to verify the feasibility of the colony based communication network, we next demonstrated that an individual sender colony with either strong or weak RBS controlling synthase production was capable of generating responses at useful scales (Figure 4). Particularly, a single colony of the strong sender was sufficient to induce response in a strong receiver strain colony at more than 1.5cm separation. This observation indicated that colony-to-colony communication at the length scale required for network design is possible.

**Figure 3:**
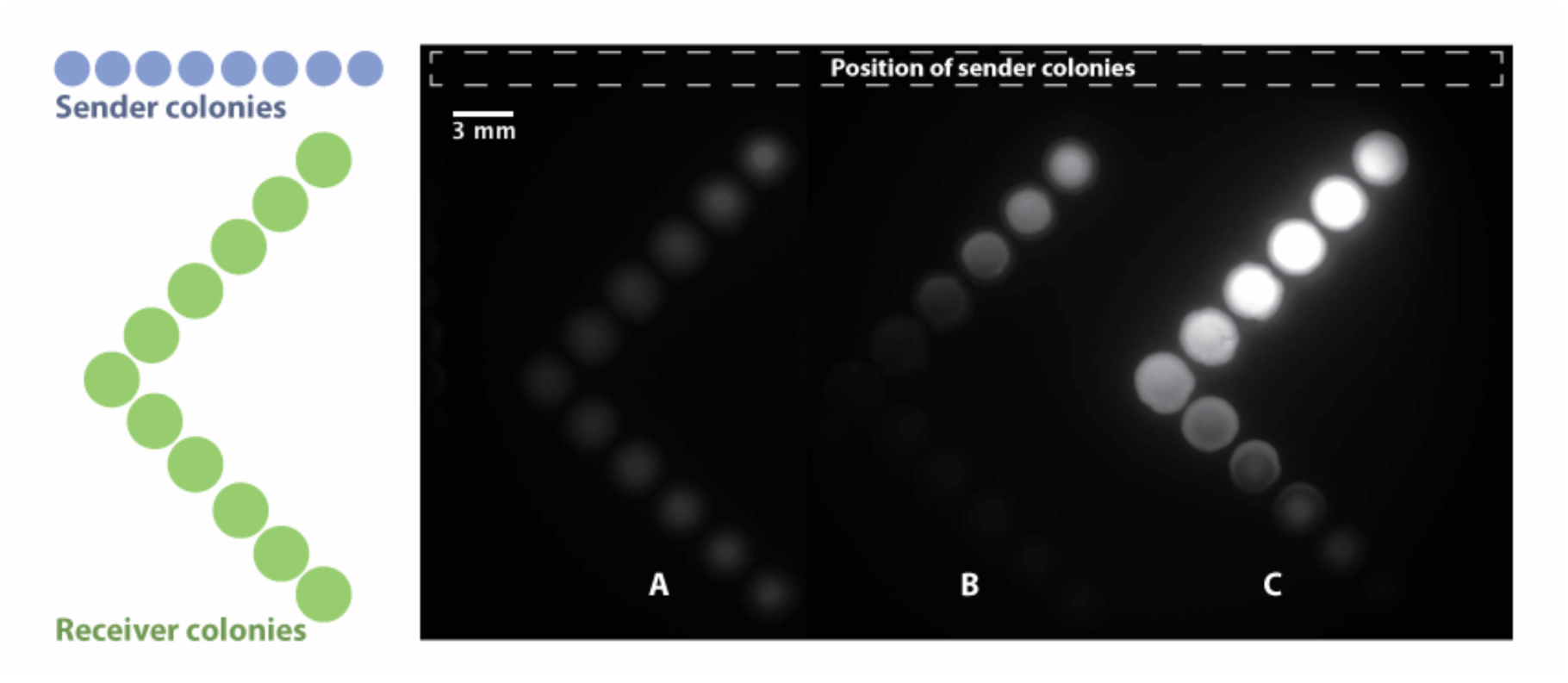
Effects of ribosome binding sites of different strength. Strong: B0034, Medium: B0032, Weak: B0033 [10] **A**. B0034 GFP, B0033 CinR **B**. B0032 GFP, B0034 CinR **C**. B0034 GFP, B0034 CinR

**Figure 4:**
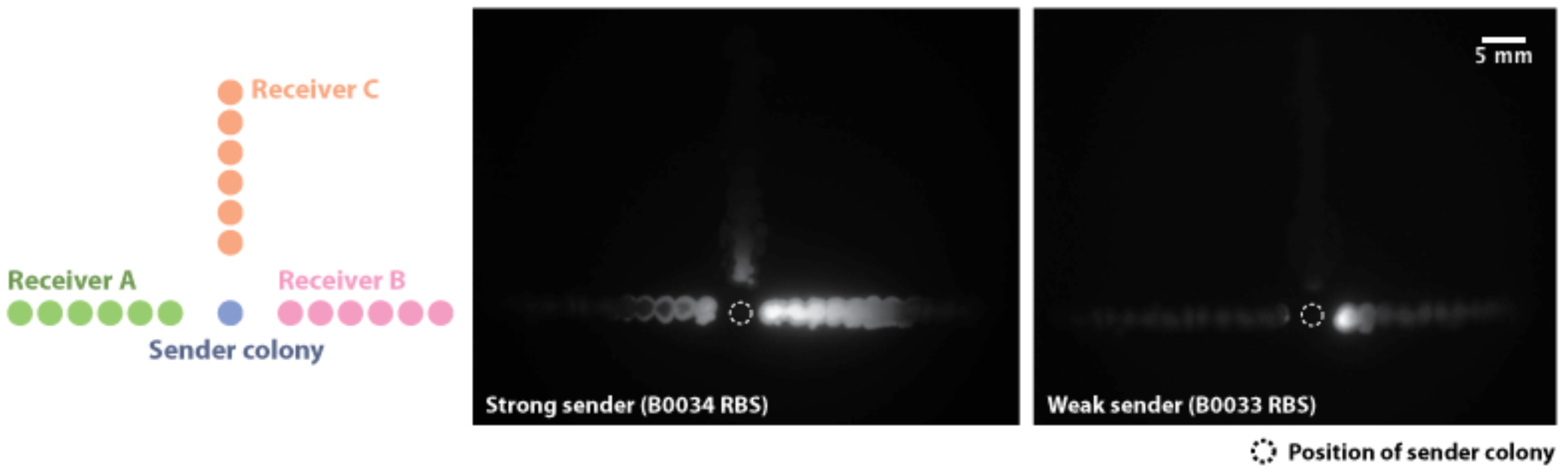
Receiver colonies responding (GFP) to single sender colonies. In each image, a sender colony was spotted between the three lines of receiver cells. **Receiver A:** B0032 GFP, B0034 CinR **Receiver B:** B0034 GFP, B0033 CinR **Receiver C:** B0034 GFP, B0033 CinR, B0034 AiiA

### Expression of AHL degradase in receiver colonies produces a signal threshold effect and response time delay

We also created reporter strains that constitutively expressed AHL degradase (AiiA) alongside the reporter circuit. Colonies of this strain start expressing measurable fluorescence at a later time point than comparable receiver without AiiA expression (Figure 5). The induction curve of AiiA expressing colonies was characterized by an initial period with no signal response, and then followed by a delayed increase comparable to that of regular receiver colonies (Figure 6). We theorize that AiiA expression initially eliminates most AHL until sufficient signals are received to saturate the enzymatic capacity of AiiA. Thus, the expression of AiiA thresholds the level of AHL and results in a signal response delay.

**Figure 5:**
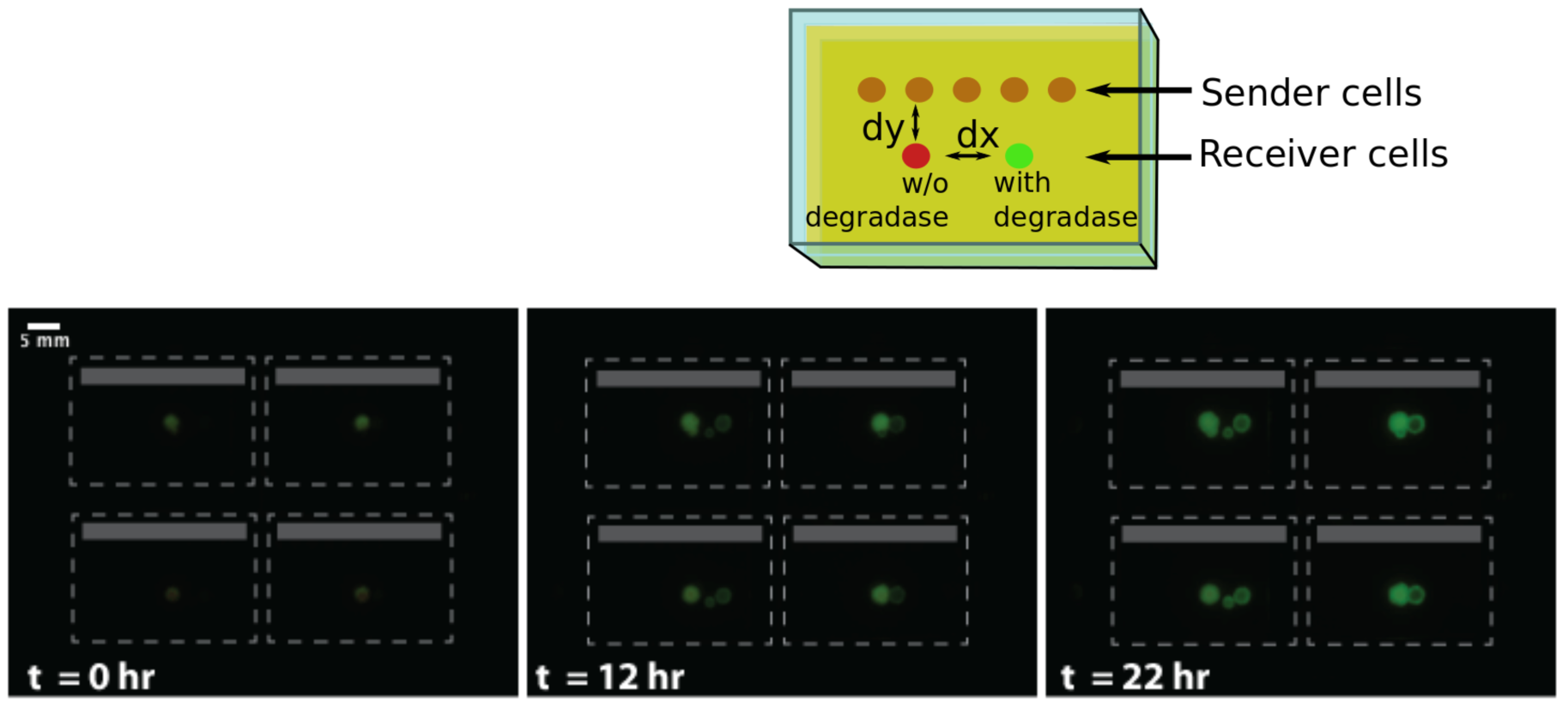
Delay of fluorescence response in receiver cells that produce degradase (right in each pair) in comparison to receiver cells without degradase (left in each pair). Four agar blocks (outlined in grey dashed lines) of the above layout are shown in these image frames. Positions of sender colonies are shaded in grey.

**Figure 6:**
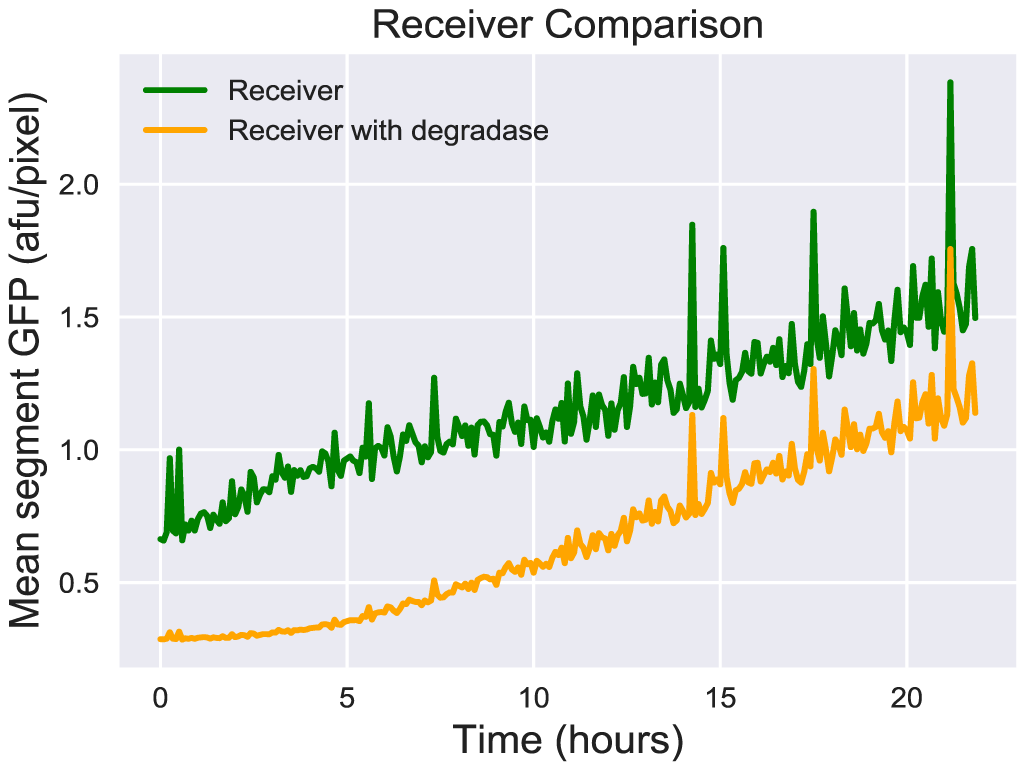
Measured fluorescence averaged over colony area. Measurements began after an overnight incubation of seeded plates. Receiver expressing AiiA started to display measurable fluorescence at a later time point.

### Receiver colonies expressing AHL degradase reduce signal response in obstructed receiver colonies

Expression of AiiA in receiver colonies has the potential of affecting signal response in neighboring colonies by diminishing the AHL level in the surrounding area of the medium. We have found that neighboring colonies were only affected in specific orientations. Regular receiver colonies appeared unaffected when placed adjacent to AiiA expressing colonies at the same distance away from the row of sender cells. This was true for colonies spotted as close as one step (2.25 mm center to center) away from the degradase-producing colonies (Figure 7). However, when a row of AiiA producing colonies was placed between the row of sender colonies and lines of regular receiver colonies, the calling distance was reduced in the regular receiver colonies (Figure 8).

**Figure 7:**
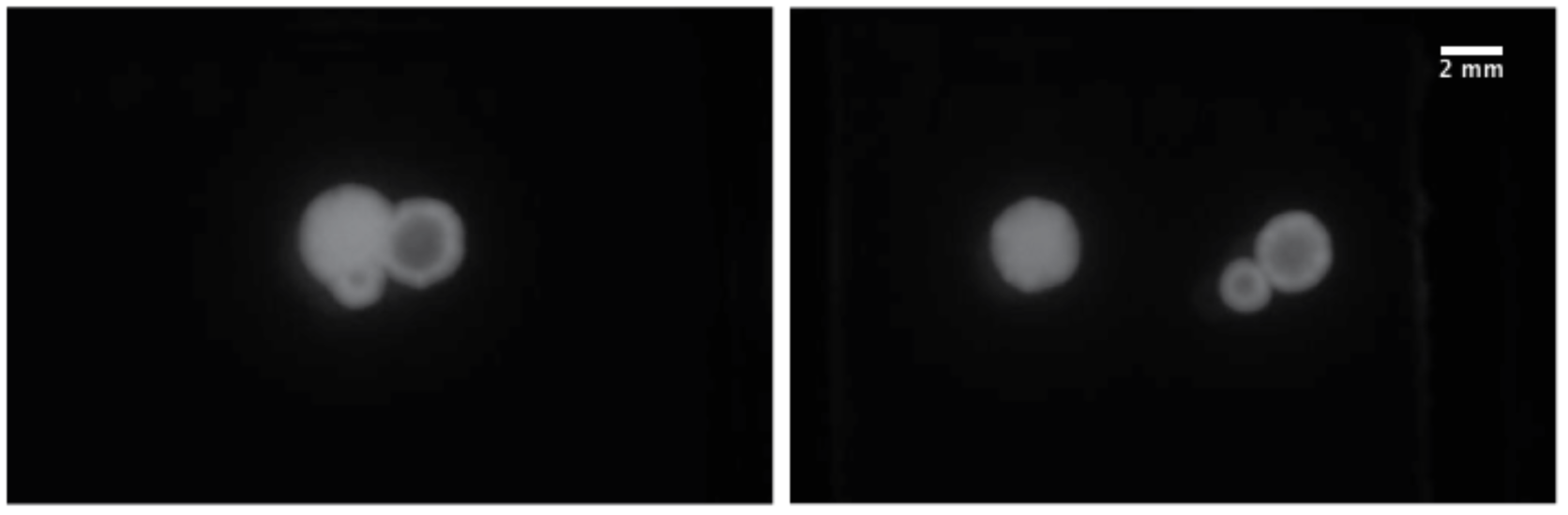
Fluorescence response in regular receiver colony (left in each pair) was constant across different proximity to degradase producing colonies (right in each pair). This figure is a close-up version of Figure 5 and follows the same experimental layout.

**Figure 8:**
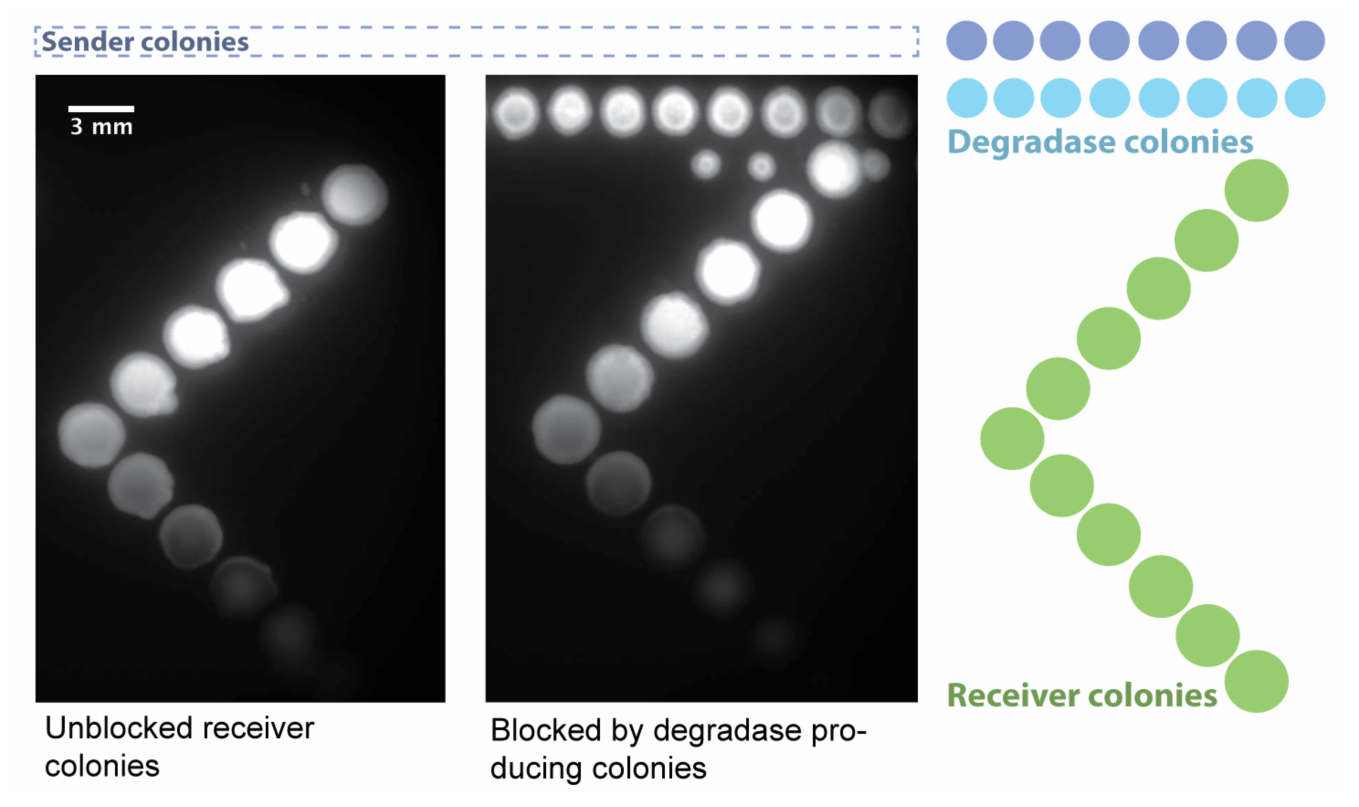
Degradase-producing colonies (top row in right image) reduced the signal response in subsequent receiver colonies (spotted diagonally). Sender colonies (not shown) were spotted as a line above the position of the degradase-producing colonies.

### A heterogeneous array of colonies propagates AHL signals for long-range communication

We investigated more complex consortia in which strains both produce and respond to AHL molecules from two orthogonal QS systems. The circuit template applied to the construction of these consortia strains was such that the presence of AHL from one QS system led to the production of the synthase corresponding to the other QS system. Figure 2 depicts the experimental demonstration we pursued for this consortium.

The array consisted of two strains occupying alternating colony positions along a line. One of the strains expressed Rhl AHL synthase upon detection of Cin AHL, while the other strain expressed Cin AHL synthase upon detection of Rhl AHL. Each colony in this system communicated with the immediately adjacent colony of a different type to propagate the signal. There were a total of six colonies in the testing array and the first colony was induced by external addition of Cin AHL. In our experiment, a colony array with center-to-center separation of 9.55mm successfully demonstrated the proposed colony-by-colony signal propagation (Figure 9).

**Figure 9:**
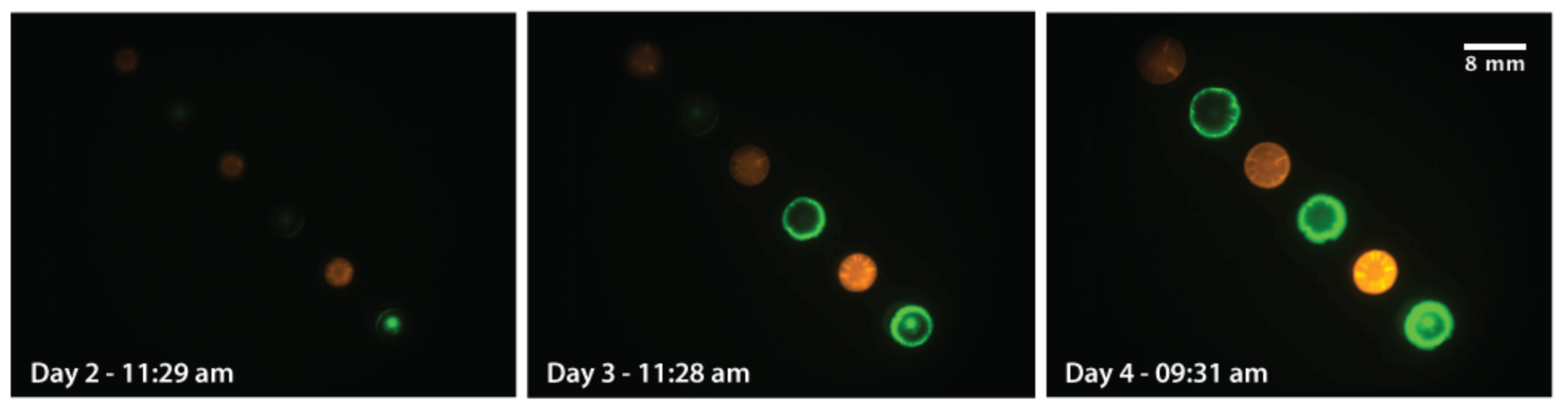
The colony array was seeded on day 0 and incubated at 37°Covernight. The signaling chain was initiated by adding 25nL of Cin AHL to the first colony at the bottom right corner of the array on day 1 around 11am. The green colonies express GFP and Rhl AHL synthase upon detection of Cin AHL, while the red colonies express RFP and Cin AHL synthase upon detection of Rhl AHL.

**Figure 10:**
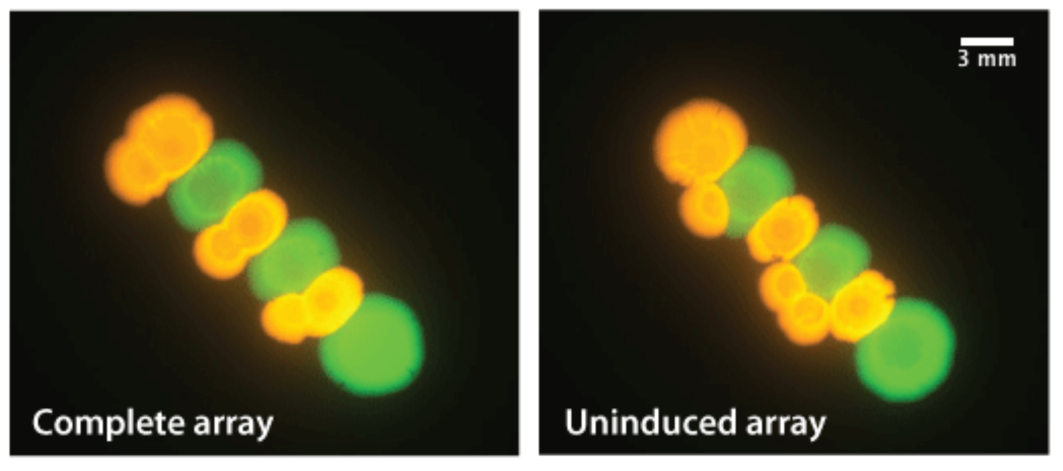
A colony array with 3.18mm center-to-center separation on day 4. Formation of an intracellular positive feedback loop among the two colony types caused self-activation of the array without external induction of the first colony.

We tested several control conditions in parallel to ensure that the communication array behaved via our intended mechanism. One significant concern was the potential of forming a positive feedback loop between adjacent colonies of different strains, leading to self-activation of the communication array. We have observed this phenomenon when the two types of colonies were in physical contact (Figure 10). In this case, the colony array responded in full without external induction of the first colony. However, this self-activation was not observed in the communication array with 9.55mm separation (Figure 11). Its uninduced control array fluoresced only at background level after four days. Furthermore, two additional control arrays showed that the signal was predominantly propagated between adjacent colonies. When one colony of either type was absent from the array, the signal response was significantly diminished beyond that gap (Figure 11). These results confirmed that the specific colony placement was crucial for the signal propagation and that the signal was propagated colony-by-colony.

**Figure 11:**
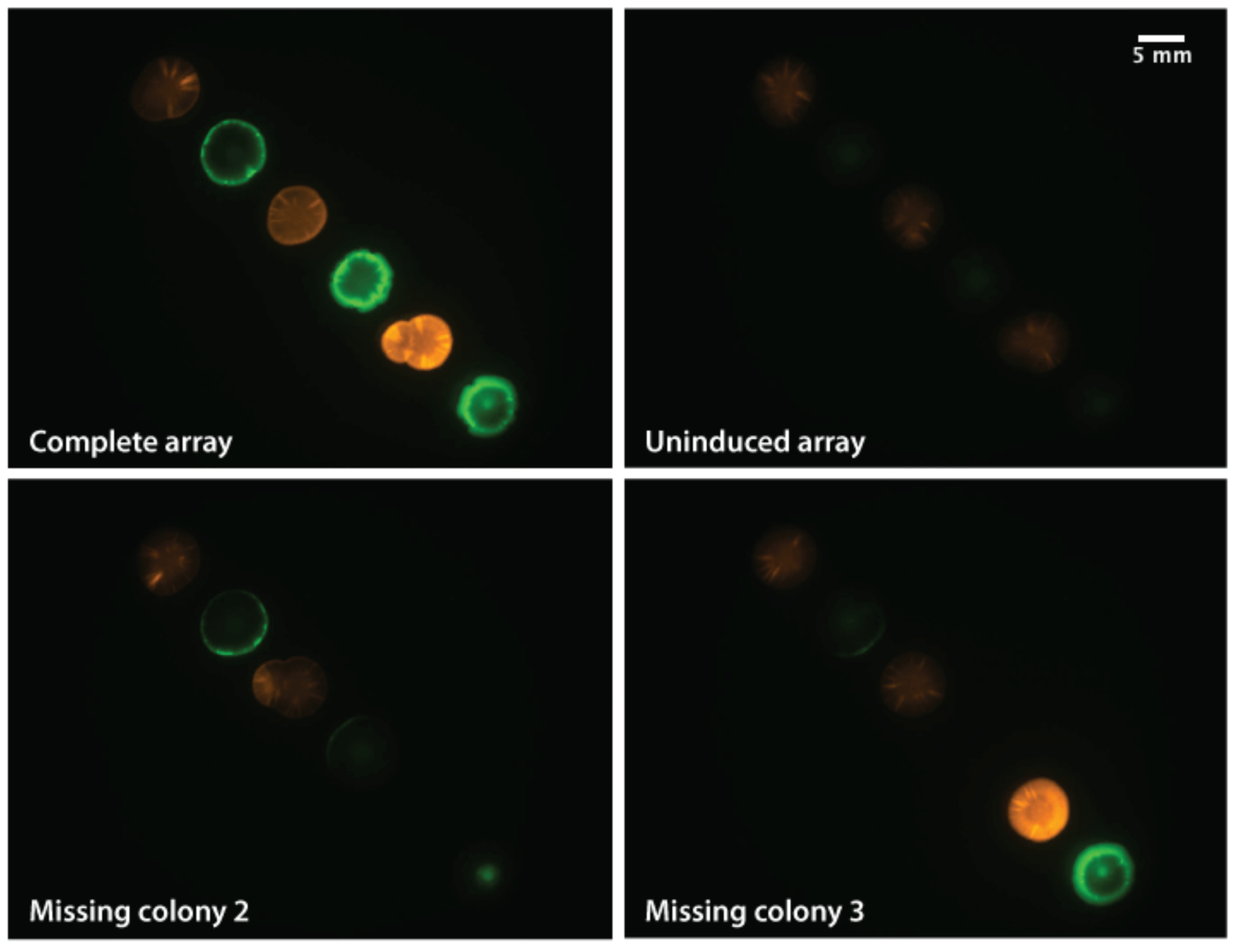
A colony array with 9.55mm center-to-center separation on day 4. The uninduced control array fluoresced only at background level. (top right image) When either the second (bottom left image) or third (bottom right image) colony was absent from the array, the signal response was significantly diminished beyond that gap.

### The heterogeneous array of colonies transmits signal response at constant rate

Image analysis has confirmed and quantified the sequential signal response observed in the array of six colonies described in the previous section. Fluorescence values were normalized and averaged over colony area detected at the specific time points. Figure 12 plots the resultant mean fluorescence values across time for each of the six colonies. These response profiles resemble induction curves. The response presumably initiated when a sufficient level of AHL signals first arrived at the colony, then the response linearly increased with the level of AHL until it reached the maximal induction, at which point the fluorescence values plateaued.

These profiles also clearly display the differences in response onset times among the array of colonies. In Figure 13, response time is approximated by the time at which a colony fluoresces at 25% of the maximal level achieved in the respective channel. All colonies have entered the inducible range of the response profile at this threshold. Figure 14 shows that response time increases linearly with colony number and that the signal transmits at a constant rate of 12.8 hours per colony or 0.075 cm per hour.

**Figure 12:**
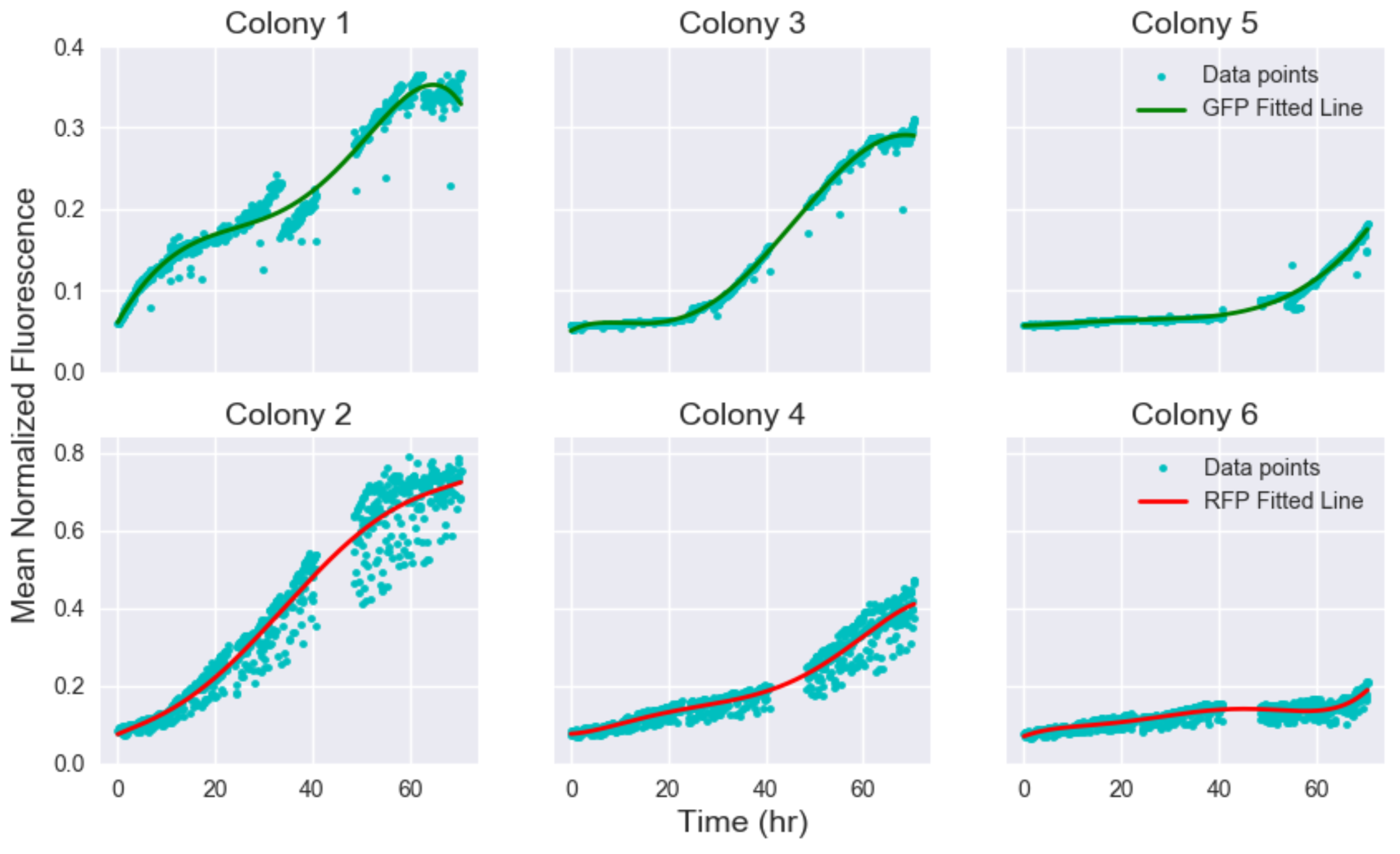
Mean normalized fluorescence over time for each of the six colonies. Fitted curves are polynomials of order 5. Signal response propagated from colony 1 to colony 6, which corresponds to left to right in this figure. Refer to Figure 2 for the patterning of the colony array.

**Figure 13:**
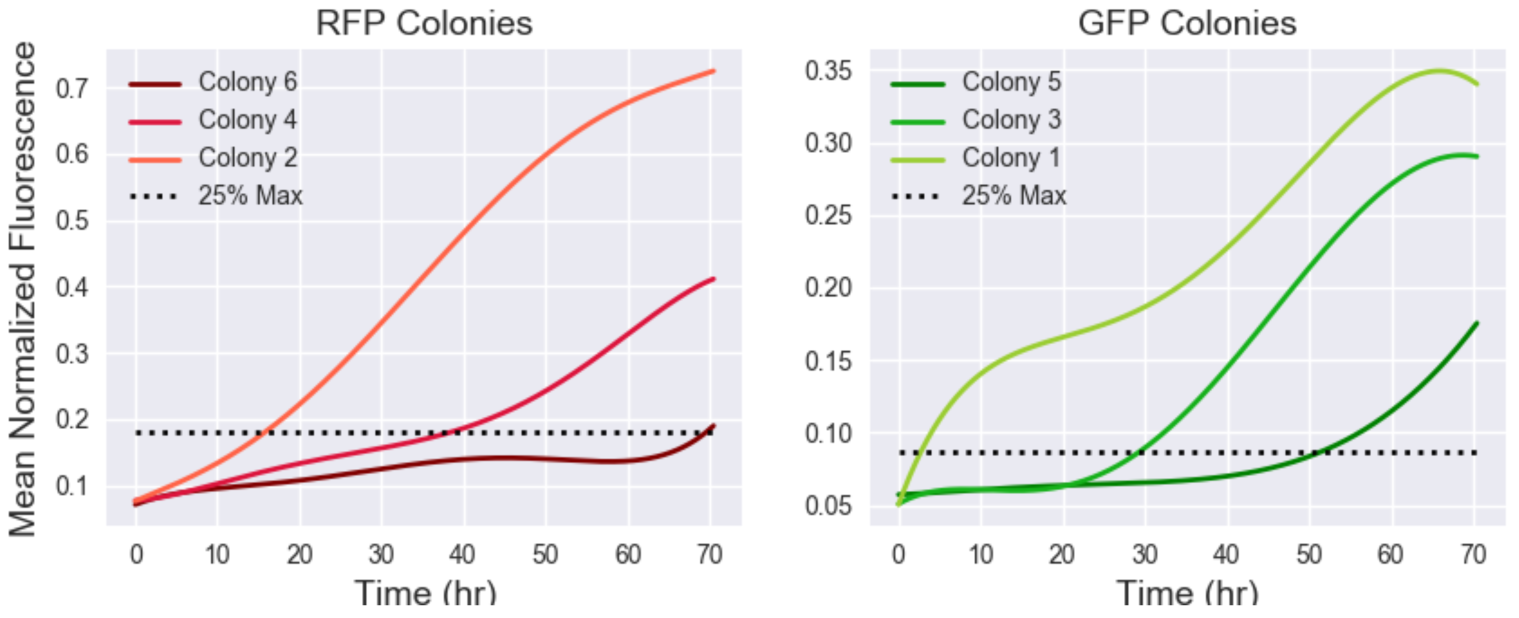
Fitted curves of mean normalized fluorescence over time in comparison. Signal response propagated from colonies 1 to 6. Refer to Figure 2 for the patterning of the colony array.

**Figure 14:**
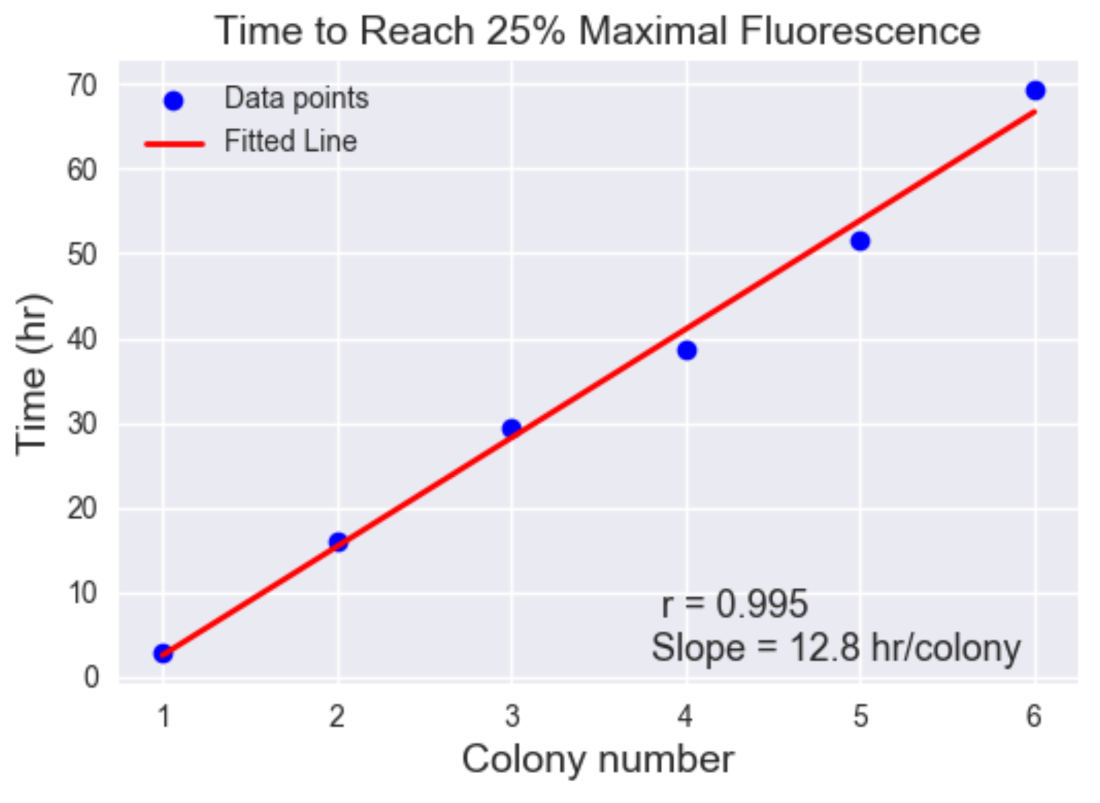
Time at which individual colonies fluoresced at 25% of the maximal level achieved in the respective channel

**Figure 15:**
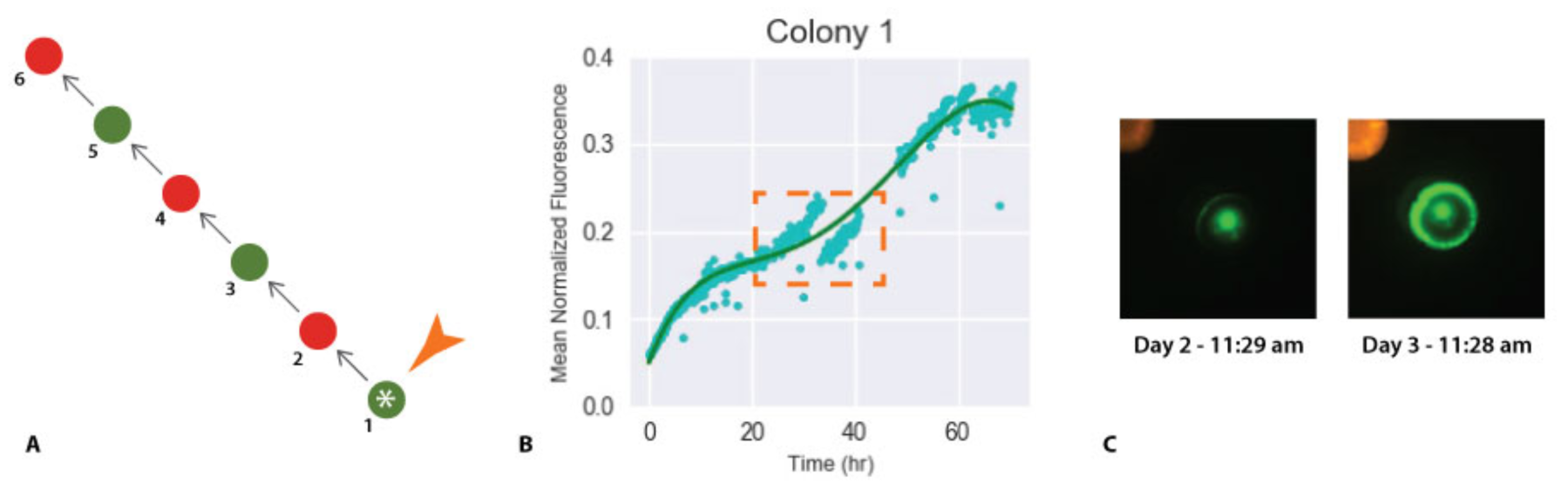
Discontinuity in the response profile of colony 1. **A** Colony 1 was the first colony in the array and its center was induced by external AHL addition. **B** The region of discontinuity in response curve. **C** The formation of inter-colony structure over time.

Several issues are evident in the data set. Firstly, some time points are missing between hour 40 and 50. Secondly, RFP expressing colonies, numbered 1, 3 and 5, produced mean fluorescence values with large spread and noise in comparison to the GFP expressing colonies because the movie exhibited a high frequency cyclic shift in detected fluorescence values in the RFP channel. Finally, the response profile of colony 1 contains an unexpected discontinuity, highlighted in Figure 15 B. The irregularity is caused by the detection and progressive formation of an inter-colony structure. In earlier time points, only the center of colony 1 fluoresced because external AHL was added to the center to initiate signal propagation. The area of this fluorescing center was detected and used for normalizing fluorescence values. In later time points, a ring structure developed at the colony edge, presumably due to feedback from the adjacent colony (Figure 15 C). Once the outer ring structure fully developed, a larger fluorescing area was detected for normalization, resulting in a sudden decrease in mean fluorescence values. We intend to address these problems in the next set of experiments by sampling a complete set of time points, replacing RFP with GFP and normalizing fluorescence values using areas detected in bright field images.

## Model

Through these experiments, we were able to quantify the length and time scales of communication of microbial consortia embedded in agar. In addition, we developed a reaction-diffusion model that captures the principal features we observed from these experiments. With this model, we can investigate how parameters impacting protein expression, cell growth, signal sensing, and signal production lead to signal cascades. The model also allows us to estimate AHL concentration as a function of time and position, which cannot be viewed directly through experiment.

The model considers nutrient-dependent cell growth, nutrient-dependent protein production, signal synthesis via synthase proteins, signal diffusion, cell diffusion, and nutrient diffusion. As the circuit is relatively simple, we elected for the model to use a discretized approximation of a continuous reaction-diffusion system. In this discretization, the agar plate is compartmentalized into chambers 1 mm wide by 1 mm long by 5mm deep, the depth of the agar in the experiments. The concentration of model species takes on a single value within each voxel. To calculate the discretized approximation of diffusion, the model calculates the rate of change in molecule concentration by applying Ficks’ First Law to the surfaces dividing adjacent rectangular chambers. For indices (i, j) and (x, y) on a grid defining chamber locations on a plate P, and chamber width w, the contribution of diffusion to the discrete approximation of the spatial derivative of a species s takes the form:

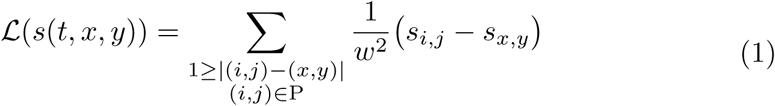

All other model equations make use of Hill functions to model nonlinear, bounded activity promoted by small molecules.

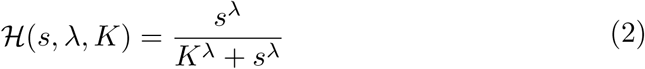

While protein production and cell growth depend on nutrient availability, only cell growth actually consumes nutrients. Each time derivative of these species involves production terms that are proportional to a production parameter, the nutrient availability in that chamber, and the concentration of the producing species (e.g. synthase is the producing species for AHL). Nutrient availability is a nonlinear function of the local nutrient concentration, defined by a Hill function [11].

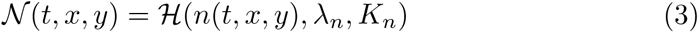

While protein production and AHL production depend on nutrient availability, only cell growth actually consumes nutrients. This feature is anchored in the assumption that the metabolic investment in the circuit’s component proteins is minor compared to all other cellular activity. In addition, we assume that protein degradation and diffusion are negligible. This assumption is supported by the observation that patterns of fluorescence within a colony are stable throughout each experiment performed. The full system of ordinary differential equations defining the model is listed here. Furthermore, the model is built with realistic parameters and units for AHL diffusion and concentration, including time, but we leave the units describing cell density, protein quantity, and nutrient concentration arbitrary [11].

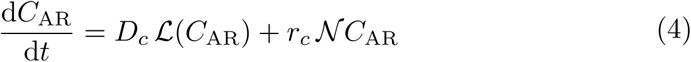

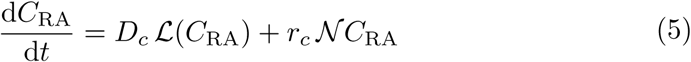

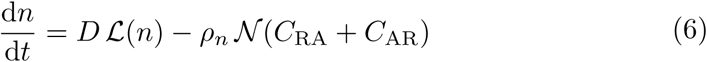

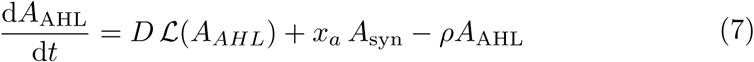

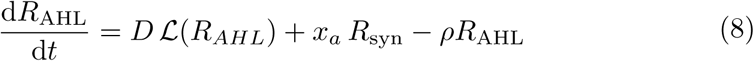

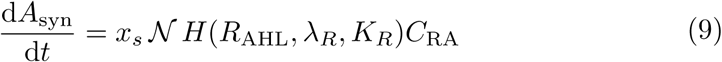

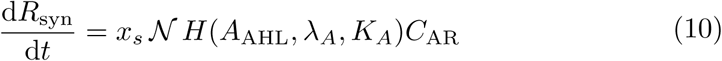

We performed simulations of each experiment and discovered, by hand, model parameters (Table 1 including initial nutrient and density values) that yield similar behavior as we observed in experiments. As was done for each experiment, the simulation allows colonies to grow for a simulated 18 hours before the addition of AHL. Below are the first and last frames of a simulation of the complete-array experiment described above, the first frame being immediately following the simulated addition of Cin AHL. In the setup of this experiment, two arrays were seeded onto the same plate, which is also the case in the simulation. We analyze only the array with larger separation between colonies.

The simulated array of colonies showed a constant signal propagation velocity, as we had found in the experiment. Simulations were also performed to verify that the signal propagation fails when a single colony is removed from the array, and that arrays with more closely-spaced colonies results in leaky activation of the signaling circuits. These simulation results confirm that our model of the circuit is valid and that it will allow for *in silico* predictions of more complicated communication circuits or colony patterns.

**Figure 16:**
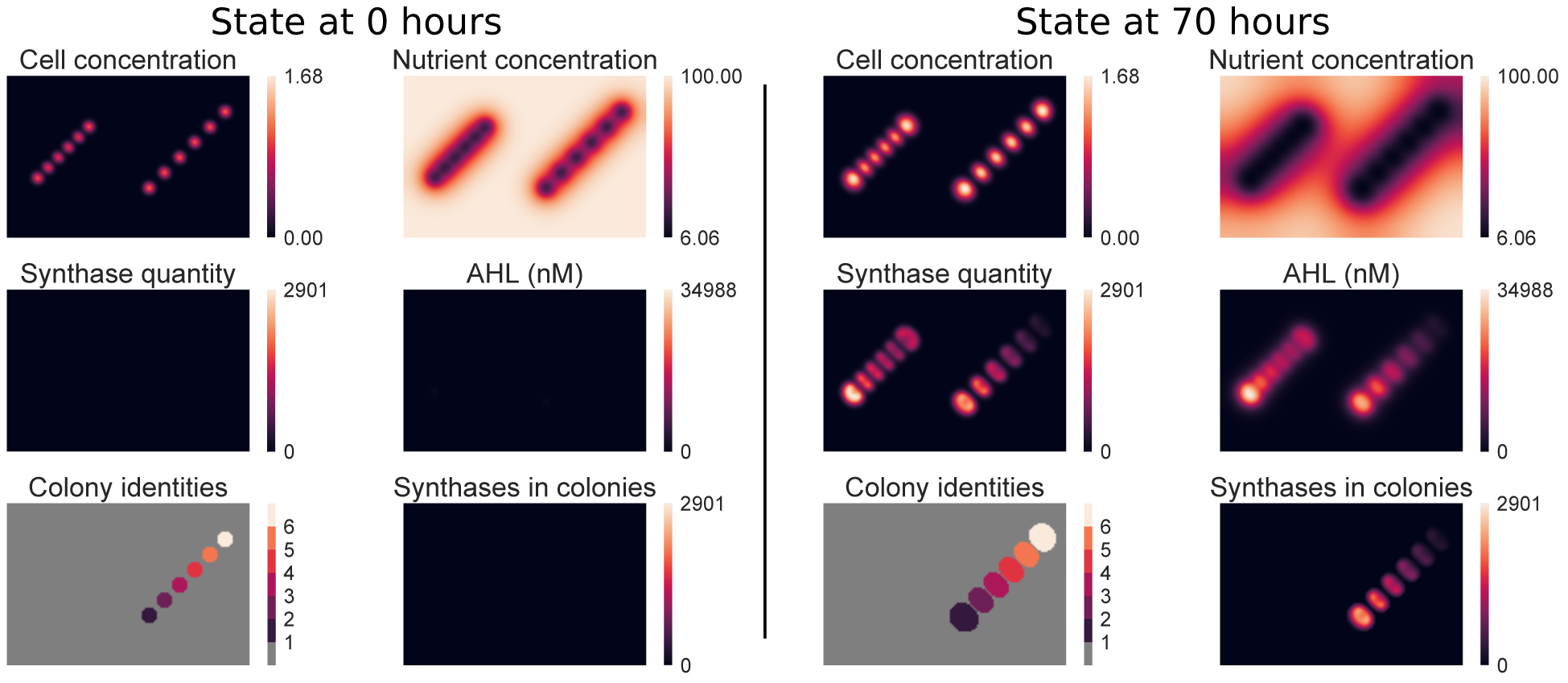
The initial and final simulation states from the simulation of the full-array cascade experiment. The bottom row of plots shows the masks that define the boundaries of the individual colonies of the signaling array of interest and the synthase distribution within those masks.

**Table 1:** Hand-picked model parameters that yield similar behavior as we observed in experiments. Parameters the character 0 in their index relate to the initial state of the simulation, 18 simulation hours before the start of the experiment. *C_aro_,Crao* are the initial values for cell density applied to the simulation chambers that correspond to the colony seed locations. *n_0_* is the initial nutrient concentration, which is uniform over the whole plate. Likewise, 2.5 mM Cin AHL is added to the seed chamber of Colony 1 following the outgrowth period.

| *Parameter* | *Value* | *Parameter* | *Value* |
| --- | --- | --- | --- |
| $D_c$ | $10^{-4} \frac{\text{mm}^2}{\text{min}}$ | $x_s$ | $20 \text{min}^{-1}$ |
| $r_c$ | $6 \cdot 10^{-4} \text{min}^{-1}$ | $\lambda_R, \lambda_A$ | 2.3 |
| $K_n$ | 80 | $K_R, K_A$ | 40nM |
| $\lambda_n$ | 2 | $\rho$ | $5 \cdot 10^{-3} \text{min}^{-1}$ |
| $D$ | $0.03 \frac{\text{mm}^2}{\text{min}}$ | $w$ | 0.75mm |
| $\rho_n$ | $3 \text{min}^{-1}$ | $n_0$ | 100 |
| $x_a$ | $0.1 \frac{\text{nM}}{\text{min}}$ | $C_{AR0}, C_{RA0}$ | 0.5 |

**Figure 17:**
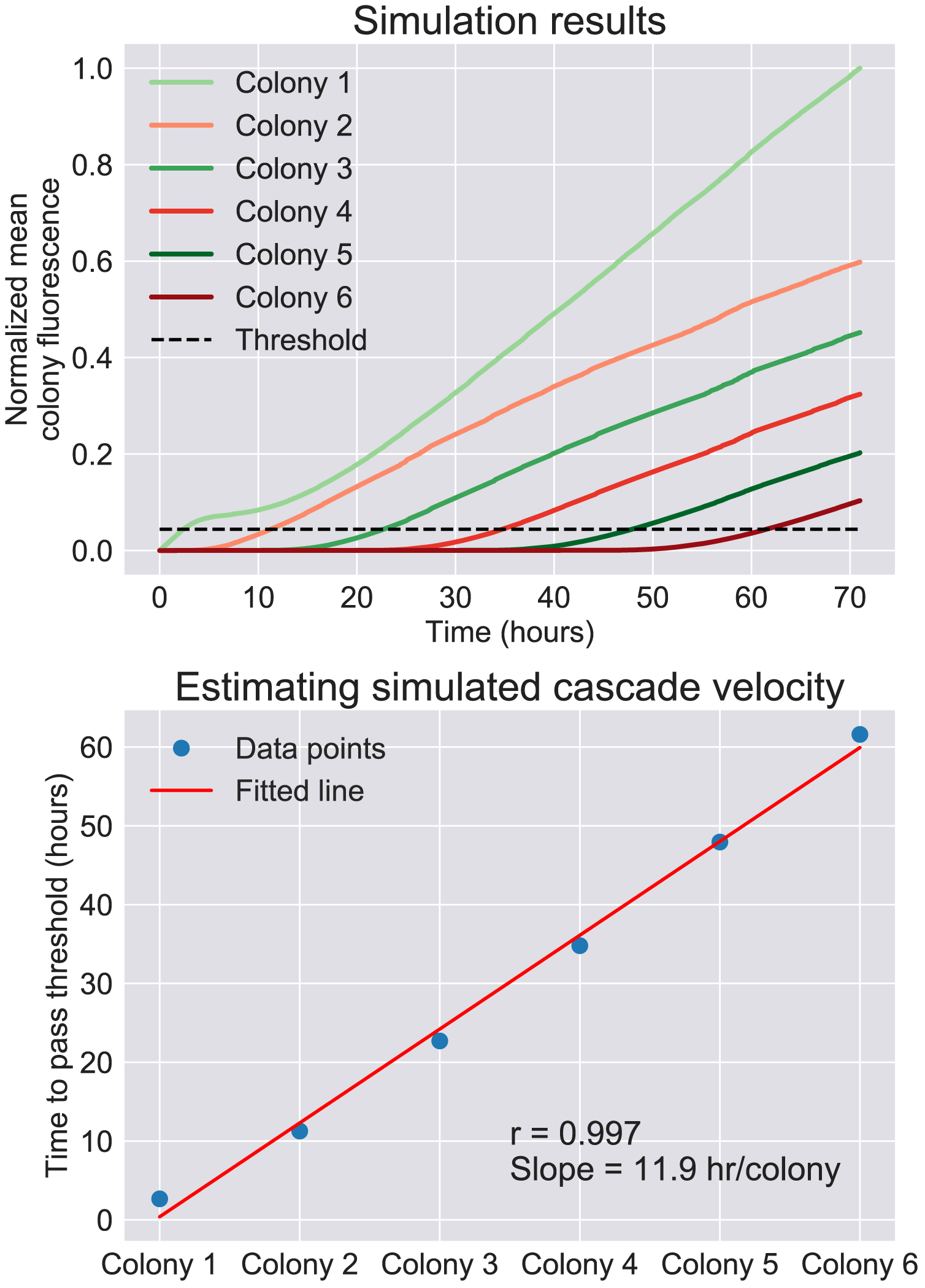
For the purposes of determining the transmission velocity of the signal cascade, we consider the synthase quantity value to represent the fluorescence from similarly-expressed reporter proteins in the experiment. The threshold for simulated data was set to the normalized colony fluorescence value that Colony 1 takes after two and a half hours, following exposure to a simulated bolus of experimenter-added AHL but before receiving AHL from its nearest neighbor. The transmission velocity in the simulation was 0.081 cm/hour, slightly faster than in the experiment.

## Discussion

In this study, we have determined several methods to tune the length and time scales of cell-cell communication in a semisolid medium and applied the range of interactions to a signal conducting colony array. Our results have shown that intentional colony arrangement can introduce new interacting behaviors and enable signal propagation. These properties can be applied to engineer sophisticated mixed communities in heterogeneous environments for advanced operations.

Parts of our observations and design correspond with those of Colisweeper, the 2013 ETH Zurich IGEM project [1]. Colisweeper is a bacterial minesweeper game in agar. The system consists of assorted colonies located in a honeycomb pattern. Individual colonies in this system employ LuxI-LuxR QS components to sense if neighboring colonies are “mines”. The distance of signal response was noted as 1.5cm in their preliminary sender-receiver diffusion characterization, which falls within the range of length scales determined in our CinI-CinR system. Colisweeper applies this response distance in a manner comparable to ours. The response distance informed the colony-to-colony separation in both colony patterns to limit signaling beyond neighboring colonies, and thus both systems incorporate intentional spatial placement in the design of multicellular systems.

Our study expands upon previous studies in two ways. First, AHL degradase is typically used for negative feedback in genetic circuits [8, 9], but here we showed that AHL degradase expression could generate signal delay and thresholding effect, as well as influence cells within the local area. These behaviors widen the range of properties that could be applied to new patterning system. For instance, AHL degradase may stabilize the off state in a bi-stable positive feedback system in semisolid medium. Second, we successfully engineered colonies capable of signal transmission, while colonies in Colisweeper are only capable of either signal emission (“mines”) or detection (“non-mines”) [1]. These transmitter colonies and the successful signal propagation demonstration confirm that it is possible to form more complex colony-based communication network. In such network, the transmitter colonies could act as cellular “wires” to connect colonies of different functions distributed throughout a certain space. An example may be a microbial consortium that self-regulates the relative population of unevenly distributed sub-communities.

Limited complexity of synthetic biological operations remains a central challenge in the field of synthetic biology, and a variety of tools and methods are under development to address this challenge [14]. Many potential solutions of this problem pertain to advancing orthogonal parts or sub-systems, both of which improve modularity in order to scale up complexity [2, 6]. In this study, we make use of multiple cell types and spatial separation in heterogeneous media to achieve partitioning. These strategies present an opportunity to perform complex biological operations in a mixed community of engineered cells living in a heterogeneous environment.

## Materials and Methods

### Circuit and strain construction

The genes used in this study were cloned by Golden Gate Assembly [10] using the constitutive promoter (J23106), ribosome binding sites (B0034, B0032, B0033), reporter coding sequences (GFP, RFP) and terminator (B0015) from the CIDAR Moclo Parts Kit [10] and hybrid QS promoters (pRhlLacO, pCinTetO) and coding sequences (AiiA, rhll, cinl) from Chen et al [8].

For the signaling circuits in the length and time scale investigation, the sender circuits expressed CinI AHL synthase under the control of Isopropyl β-D-1-thiogalactopyranoside (IPTG), which was added to the LB agar plates. The receiver circuits expressed CinR transcription factor constitutively and GFP under the control of a Cin activated promoter. The constitutive AHL degradase gene (AiiA) was located on a separate plasmid and co-transformed with the receiver circuit for the degradase-expressing receiver strain. DH5α-z1 and MG1655-z1 were used for the sender strains, while regular DH5α and MG1655 were used for receiver strains.

The signaling circuits in the communication array each consist of two genes that were cloned to the same plasmid by Gibson Assembly. One circuit expresses Rhll and GFP from separate Cin activated promoter, and another expresses CinI and RFP from separate Rhl activated promoter. Each circuit was transformed into CY026 strain, which expresses CinR and RhIR transcription factors constitutively from the genome [8].

### Plating and stereoscope recording

The spatial patterns were planned in an Excel template according to the 1536-plate grid. Seeding cultures were diluted from outgrowths of overnight LB cultures to 0.5 or 0.35 OD600. 25 nL of liquid culture was seeded on LB agar OmniTrays for each designated position using the Echo 525 Liquid Handler. Additionally, in the signal delay experiment shown in Figure 5, 1-well wide stripes of agar between the colony pairs were physically cut out and removed immediately after seeding to form isolated blocks of agar medium for each colony pairs. All seeded LB agar OmniTrays were incubated overnight at 37°C. 25 nL of 2.5mM Cin AHL inducer was added on colony 1 of the communication array the subsequent morning using the Echo 525 Liquid Handler. Agar plates were stored at room temperature thereafter. Images of the plates were taken using an Olympus MVX10 microscope in RFP and GFP channels every 5 to 12 hours. Movies of a subset of plates were taken at 5 min interval for 12 to 70 hours.

## Acknowledgments

We thank Andrey Shur for technical assistance with the microscope, Shan Huang for sharing *E. coli* stock and other Murray Lab members for useful discussions. Joy Doong was supported by the Amgen Scholar Program and James M. Parkin is supported by the Institute for Collaborative Biotechnologies through grant W911NF-09-0001 from the U.S. Army Research Office. The content of the information does not necessarily reflect the position or the policy of the Government, and no official endorsement should be inferred. Plasmid vectors and non-coding regions were provided by Douglas Densmore at the Cross-disciplinary Integration of Design Automation Research lab (Addgene Kit # 1000000059). Quorum sensing promoters, coding sequences and CY026 strain were provided by Matthew Bennett (Addgene Plasmid # 65954, 65952, 72340).

